# Responsiveness of cerebral cortex to hippocampal inputs depends on brain region and brain state

**DOI:** 10.64898/2026.08.13.744702

**Authors:** Faycal Rezaig, Wyna Gagliano, Guillermo Lazcano, Pablo Fuentealba, Alain Destexhe

## Abstract

Gamma oscillations (30–90 Hz) are a prominent signature of cortical network state, but whether they facilitate or hinder inter-areal communication remains unresolved. The communication-through-coherence hypothesis posits that gamma enhances transmission between areas, whereas recent computational work suggests that high-amplitude gamma oscillations may instead filter incoming inputs and reduce their impact. To distinguish between these accounts, we used a well-defined physiological input—sharp-wave ripple (SWR) complexes—to probe cortical responsiveness via two parallel monosynaptic pathways: from ventral CA1 to prefrontal cortex (PFC), and from dorsal CA1 to retrosplenial cortex (RSC). By classifying the cortical state immediately preceding each ripple as low- or high-amplitude gamma, we found that PFC responses to ripples were significantly larger during low-amplitude gamma states, an effect carried by the ventral CA1–PFC pathway and driven primarily by ripples during quiet wakefulness. RSC showed no such state-dependent modulation. A mean-field model of PFC reproduced the enhanced responsiveness during low-amplitude gamma and revealed that this modulation depends on the excitatory–inhibitory balance of the afferent input and on the level of recurrent excitation—providing a mechanistic explanation for the distinct behaviors of PFC and RSC, which differ in their local recurrent connectivity. Extending the model to a chain of cortical areas predicted that low-amplitude gamma supports robust propagation of activity across regions, whereas high-amplitude gamma confines it locally. Together, these results argue that low-amplitude gamma, rather than strong gamma synchronization, constitutes a favorable substrate for communication between brain areas.

## 1. Introduction

The response of a neural circuit to an input depends not only on the properties of the input itself, but also on the state of the circuit at the moment the input arrives (Hasenstaub et al., 2007; Fries, 2015). In the cerebral cortex, gamma-band activity (30–90 Hz) provides a readout of this ongoing state. Gamma oscillations are not merely a spectral marker, because they emerge from the interaction between excitatory pyramidal cells and fast-spiking inhibitory interneurons (Cardin et al., 2009; Sohal et al., 2009; Buzsáki and Wang, 2012), the amplitude and frequency of oscillation are modulated by recurrent excitation-inhibition interactions, and its instantaneous amplitude reflects the momentary balance between excitation and inhibition of the local network (Buzsáki and Wang, 2012).

Gamma waves have been implicated in a broad range of cognitive functions, including attention, memory, and sensory processing (Tallon-Baudry, 2009; Köster et al., 2022). Mechanistically, gamma oscillations have been proposed to regulate information flow in cortical networks by modulating neural gain and coordinating spike timing across populations (Fries, 2005; Buzsáki and Wang, 2012; Bastos et al., 2012). Yet their precise functional role — and whether they enhance or impede communication across brain regions — remains debated. According to the communication-through-coherence framework, gamma synchronization increases the gain of communication between cortical areas (Fries, 2005, 2015). In contrast, computational modeling has suggested that strong gamma oscillations reflect internally generated activity that filters incoming inputs, thereby reducing rather than increasing their impact, independently of the mechanism that generates the oscillation (Susin and Destexhe, 2021). These two views make opposite predictions about how a cortical circuit engaged in gamma should respond to an external input, and discriminating between them requires a well-defined physiological input delivered while the ongoing gamma state is monitored. Hippocampal sharp-wave ripples (SWRs) provide such an input: they are natural, phasic excitatory volleys that propagate to the cortex and can be detected reliably. Two networks receiving the same afferent volley may therefore respond very differently depending on whether they are, at that instant, in a high- or low-amplitude gamma regime. This motivates using pre-input gamma amplitude as a readout of cortical state, and raises the question of whether the excitatory–inhibitory balance that shapes gamma also governs how strongly an incoming input is transmitted. Here we ask whether this state-dependent filtering is shaped by local recurrent connectivity, comparing two cortical areas — PFC and RSC — that differ in their circuit architecture.

SWRs are among the most prominent population events in the mammalian hippocampus. They comprise a sharp wave, which largely reflects CA3 input, together with a fast ripple oscillation generated mainly in CA1, and occur during quiet wakefulness and NREM sleep (Buzsáki, 2015). SWRs are associated with the reactivation of waking ensemble activity (Wilson and McNaughton, 1994) on a compressed timescale (Lee and Wilson, 2002), and their influence extends beyond the hippocampus to cortical and subcortical regions (Logothetis et al., 2012; Ramirez-Villegas et al., 2015). Consistent with a role in memory processing, selective suppression of ripples impairs subsequent spatial memory (Girardeau et al., 2009), hippocampal and cortical activities are coordinated around SWRs (Sirota et al., 2003; Ji and Wilson, 2007), and ripple–cortical coupling strengthens with learning (Khodagholy et al., 2017). Together, these findings support a role for hippocampo-cortical communication during SWRs in systems consolidation.

It nonetheless remains unclear how efficiently individual ripples are transmitted to the cortex, and which factors set the strength of this transmission. Because SWRs occur almost exclusively during quiet wakefulness and NREM sleep—when cortical activity fluctuates between relatively silent and active periods—the same hippocampal event can reach the cortex in very different states from one ripple to the next. If hippocampo-cortical transfer contributes to memory consolidation, it therefore becomes important to determine whether the cortical state at the time of a ripple affects its transmission. As a measure of this state, and for the reasons above, we focused on the amplitude of ongoing gamma activity immediately before each ripple. We treat the ripple as a feedforward input to the cortex for the purposes of this analysis, while noting that hippocampo-cortical coupling is bidirectional and that the cortical state can itself bias ripple occurrence (Sirota et al., 2003).

The prefrontal cortex (PFC) is a natural target for this question. The hippocampus connects to PFC through multiple routes: a direct monosynaptic projection arises from ventral CA1 and ventral subiculum (Jay and Witter, 1991), whereas dorsal CA1 reaches PFC mainly through polysynaptic pathways relayed by structures such as CA3, the ventral hippocampus, thalamus and lateral septum. Thus, the vCA1–PFC pathway provides a well-characterized direct excitatory input to this cortical area. Functionally, prefrontal neurons are phase-locked to hippocampal rhythms (Siapas et al., 2005), and prefrontal ensembles replay learning-related activity during ripples (Peyrache et al., 2009). The CA1–PFC circuit therefore offers an ideal system to test whether the cortical gamma state before a ripple shapes the evoked response.

To determine whether this relationship is general or depends on local circuit architecture, we compared PFC with the retrosplenial cortex (RSC). Like PFC, RSC receives a direct monosynaptic projection from the hippocampus—in this case, from dorsal CA1—but with a crucial difference: this projection is predominantly inhibitory, arising from a specialized population of CA1 GABAergic neurons that innervate superficial layers of RSC (Brennan et al., 2020; Opalka et al., 2020). Beyond this difference in afferent sign, the two target areas also differ markedly in their local circuit organization. PFC pyramidal neurons form a densely interconnected recurrent excitatory network (Wang et al., 2006; Anastasiades and Carter, 2021). In contrast, the superficial RSC contains little recurrent excitation, strong inhibition, and a population of hyperexcitable, weakly adapting principal neurons (Brennan et al., 2020). Moreover, ripple-associated activity activates RSC interneurons while suppressing excitatory neurons, consistent with a feed-forward inhibitory response (Opalka et al., 2020). Comparing these two areas—both receiving monosynaptic hippocampal inputs, but differing in afferent sign and local architecture—allows us to test whether the link between cortical state and responsiveness is shared across the cortex or is instead shaped by local circuitry.

To complement the experiments and gain mechanistic insight, we developed a biophysically constrained mean-field model of coupled excitatory and inhibitory adaptive-exponential populations (Zerlaut et al., 2018; di Volo et al., 2019). This modeling approach enables us to systematically dissect the factors that determine responsiveness—such as the excitatory–inhibitory balance of the afferent input and the strength of recurrent excitation—which would be difficult to isolate experimentally. Moreover, by extending the model to a chain of interconnected cortical areas, we could test whether the observed differences in responsiveness translate into differences in inter-areal propagation.

We found that the cortical response to hippocampal ripples was larger during low-amplitude gamma states and smaller during high-amplitude gamma states in PFC, whereas no such state-dependent modulation was observed in RSC. To our knowledge, this provides the first direct experimental evidence, in PFC, that gamma oscillations exert a bidirectional, amplitude-dependent effect on input transmission: low-amplitude gamma favors responsiveness, whereas high-amplitude gamma suppresses it. This finding is consistent with the filtering account (Susin and Destexhe, 2021) and challenges the simplest interpretation of communication-through-coherence, which would predict enhanced transmission during high-amplitude gamma synchronization. Our model reproduced the two gamma states and captured the stronger ripple response in the low-gamma state; simulations further indicated that this modulation depends on the excitatory–inhibitory balance of the incoming input, and that the area difference between PFC and RSC can be accounted for by their distinct recurrent connectivity and afferent bias. Extending the model to a chain of cortical areas showed that low-gamma states support more reliable inter-areal propagation, whereas high-gamma confines activity locally. Overall, our results suggest that ongoing gamma amplitude is inversely related to the responsiveness of a cortical circuit to incoming inputs, and highlight the excitatory–inhibitory balance of the afferent drive and local recurrent connectivity as key determinants of this relationship.

## 2. Methods

### 2.1 Experimental recordings

In rats the local field potentials and the activity of individual neurons were recorded in the prefrontal cortex and the retrosplenial cortex at both the dorsal and ventral sites as well as in the hippocampal CA1 area (the dorsal part being CA1d and the ventral part CA1v) while the animals were in quiet wakefulness (QW) and during NREM sleep. In the CA1 area ripples were identified and the cortical response was then synchronized to the onset of the ripples. The state of the cortex before the ripples was classified as either low or high-amplitude gamma according to the gamma power measured in a time window that lay one second before the onset of the ripples.

### 2.1 Animals, surgery and recordings

Experiments were performed on 17 adult male Sprague-Dawley rats (postnatal days 40-60, 300-350 g), were obtained from the Center for Innovation on Biomedical Experimental Models (CIBEM, https://cibem.bio.puc.cl/) at the Pontificia Universidad Católica de Chile. Rats were housed under a 12 h light/dark cycle with food and water available ad libitum, and habituated to handling and to the recording environment before implantation. All procedures were approved by the institutional ethical committee and are reported in accordance with the ARRIVE guidelines (https://arriveguidelines.org).

Under isoflurane anaesthesia (4 % induction, 1.5-2 % maintenance) in a stereotaxic frame, 50 um diameter tungsten tapered-tip electrodes mounted on a custom 3D-printed microdrive were implanted in the right hemisphere in dorsal CA1 (CA1d), ventral CA1 (CA1v), retrosplenial cortex (RSC) and prefrontal cortex (PFC); ground and reference screws were placed in the skull anterior to bregma. Animals recovered for at least one week before recordings. Signals were acquired with an Intan RHD system at 20 kHz, in 60-90 min sessions, with synchronised video. Electrode positions were verified post hoc by electrolytic lesions and Nissl staining.

### 2.2 Brain-state scoring, ripple detection and spike sorting

The brain states were visually assessed in 10-second intervals based on the CA1d and PFC field potential, its spectrogram, and the video record; quiet wakefulness was defined as a state involving mixed high-frequency activity together with little or no movement, NREM sleep as prolonged low-frequency (0.1–4 Hz) cortical activity in the absence of movement, and REM sleep as being characterised by prominent hippocampal theta activity in the absence of movement; nevertheless, the REM episodes were excluded from the present analysis. The ripples were identified by applying band-pass filtering to the CA1 signal in the ripple frequency range (100-250 Hz, with zero-phase characteristics), then rectifying it, low-pass filtering the envelope, z-scoring it and keeping only those events that lasted at least 50 ms, the onset and offset of the events being determined from crossings of a lower threshold and with a refractory period introduced in order to prevent duplicate detections; the detections were checked visually. The spikes were obtained from the 600 to 5000 Hz band, sorted using Kilosort2 and then refined manually; clusters which could not be clearly separated were considered to be multi-unit activity.

### 2.3 Pre-ripple gamma classification and response quantification

To define the pre-event cortical state for each detected hippocampal ripple, 1/*f* frequency-weighted LFP gamma power was quantified within a 1.0 s window ending 100 ms prior to ripple onset (spanning from −1.1 to −0.1 s relative to onset). This 100-ms buffer zone was implemented to prevent spectral leakage from the ripple from contaminating the pre-ripple cortical state estimates. Gamma frequency bands were established according to regional spectral profiles: 40-80 Hz for PFC and 40-60 Hz for RSC. Following Median Absolute Deviation (MAD) outlier rejection, events were within-subject *Z*-score normalized and classified into tertiles, defining the first tertile as the Low pre-ripple gamma condition and the third tertile as the High pre-ripple gamma condition.

Cortical neuronal activity, comprising single-unit activity (SUA) and multi-unit activity (MUA), was binned at 10 ms and aligned to ripple onset. To isolate the net neuronal response exclusively driven by the ripple from background network activity, real event responses were baseline cross-standardized against shuffled controls. For each real event, 100 shuffled timestamps were generated by randomly jittering real ripple timestamps by ±2 to 10 s within the same behavioral state epoch, breaking precise temporal coupling to the ripple while preserving local firing rates. Net evoked reactivity was computed for each unit as:

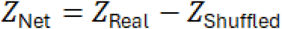

Net evoked responses between Low and High pre-ripple gamma conditions were compared across units for each 10 ms time bin within a window spanning from −1.1 to 1.1 s relative to ripple onset using a two-tailed Wilcoxon signed-rank test, with p-values adjusted for multiple comparisons using the Benjamini-Hochberg FDR procedure (α= 0.05). Significant epochs are indicated above each trace.

### 2.4 Mean-field model: formalism

The cortical network was described by a first-order reduction of the Markovian mean-field formalism (El Boustani and Destexhe, 2009; Zerlaut et al., 2018; di Volo et al., 2019), following the gamma model of Tahvili and Destexhe (2024) from which the present implementation is directly derived. The state variables are the mean firing rates of the excitatory (*e*, regular-spiking) and inhibitory (*i*, fast-spiking) populations, evolving on a Markovian time bin *T*:

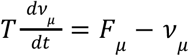

Finite-size covariance terms of the second-order formalism were not integrated, as in the reference gamma model: stochastic fluctuations are supplied by an Ornstein–Uhlenbeck process added to the external input (Section 2.5), and gamma oscillations are generated deterministically by the transmission-delay asymmetry (Eq. 10). Each population is a network of adaptive exponential integrate-and-fire neurons (Brette and Gerstner, 2005):

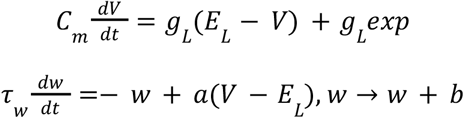

Adaptation is restricted to the excitatory population; for the fast-spiking population, *w* = 0 at all times.

The function F gives, as a function of the rates of excitatory and inhibitory input, the steady-state output rate of a population. It is derived semi-analytically from the first two statistical moments of the membrane potential, the conductances and the subthreshold moments being:

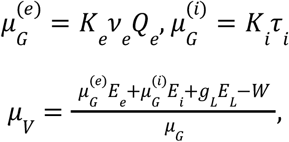

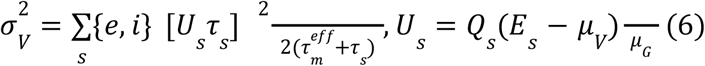

and the output rate is expressed through an effective, fluctuation-dependent threshold:

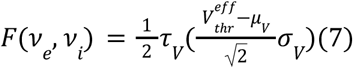

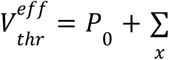

*With the normalized effective time constant*

Each variable is centred and scaled by fixed normalization constants 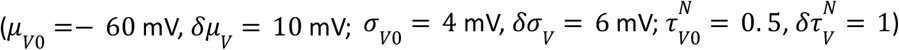, and *delimits the fluctuation-driven regime*.

The coefficients P per cell type are those of the reference implementation of di Volo et al. (2019, ModelDB 263236), used without refitting. Spike-frequency adaptation of the excitatory population enters equation (5) through its stationary value:

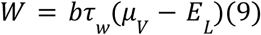

Gamma oscillations arise when synaptic transmission delays are taken into account, *F*_*µ*_ in equation (1) being evaluated at delayed rates. Following Tahvili and Destexhe (2024), the oscillation requires an asymmetry between transmission delays, implemented as a multiplicative factor *ε* on a base delay *τ* _*l*_ :

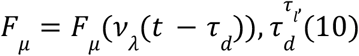

where same-population pathways (*e* →, *i* →) carry the longer delay and cross-population pathways (*e* →, *i* →) the shorter one, as in the reference model. With *τ* _*l*_ = 3 ms, *ε* = 0. 05 and an integration step *dt* = 0. 1 ms, the delays quantize to 3.1 and 2.8 ms respectively (delay difference 0.3 ms). Simulations used *dt* = 0. 1 ms throughout

### 2.5 Model parameters and gamma states

The cortex was modelled using a two-population mean-field model based on adaptive exponential integrate-and-fire neurons, namely the regular-spiking excitatory (RS) and fast-spiking inhibitory (FS), in accordance with the semi-analytical transfer-function formalism, and included spike-frequency adaptation for the RS population (Zerlaut et al., 2018; di Volo et al., 2019). Gamma oscillations were achieved by introducing an asymmetry in the synaptic transmission delays between the populations (Tahvili and Destexhe, 2024).

The low and high amplitude gamma states were obtained by changing the effective leak reversal potentials of the two groups. For the low state E_le was −68 mV and E_li was −58 mV, and for the high state E_le was −58 mV and E_li was −72 mV. These operating points were not merely assumed; instead, E_le and E_li were varied over a grid ranging from −76 to −54 mV in 2 mV increments, giving a total of 144 points, and each point was characterised by gamma peak frequency, the power in the 30–90 Hz band, the baseline RS/FS rates and viability, after which the two points were chosen. In the following, the term “low or high-gamma” refers to the gamma power, with the low-gamma state having the higher peak frequency.

**Table 1:** Model parameters.

| Parameter name | Value |
| --- | --- |
| $G_l$ (leak conductance) | 10 nS |
| $C_m$ (capacitance) | 200 pF |
| $E_l$ (resting potential) | -65 mV |
| $E_e / E_i$ (reversal) | 0 / -80 mV |

Synaptic
| Parameter | Value |
| --- | --- |
| $Q_e / Q_i$ (quantal conductances) | 1.25 / 5 nS |
| $\tau_e = \tau_i$ (synaptic time constants) | 5 ms |
| $\tau_l$ (transmission delay) | 3 ms |
| T (mean-field time constant) | 5 ms |

RS adaptation
| Parameter | Value |
| --- | --- |
| $\tau_w$ (adaptation time constant) | 500 ms |
| $b_{RS}$ (adaptation current increment) | 60 pA |
| $a$ (adaptation conductance) | 4 nS |

Network
| Parameter | Value |
| --- | --- |
| $g_{ei}$ (inhibitory fraction) | 0.20 |
| N (total neurons) | 10,000 |
| p (connectivity probability) | 0.05 |

## 3. Results

We start by showing the experimental results characterizing the response to hippocampal inputs as a function of the gamma state, then we describe a computational model aimed at reproducing these results.

### 3.1 Experiments

The experimental paradigm consisted of simultaneous recordings in two different monosynaptically-connected pathways, the CA1 region of hippocampus, dorsal and ventral, and their respective connections in retrosplenial cortex (RSC) and prefrontal corfex (PFC). The hippocampal recordings were used to detect the occurrence of sharp-wave ripple (SWR) complexes, and the simultaneous cortical recordings were used to quantify the cortical response to the SWR input, in the two areas. We divided the cortical response according to the cortical state prior to the stimulus, separating in low-amplitude and high-amplitude gamma oscillation states (see Methods).

The main finding, as shown in Fig. 1A, is that the PFC shows an overall larger responsiveness when the cortex displays low-amplitude oscillations, while the RSC does not show any significant difference. This augmentation of responsiveness is largely due to the SWR occurring in quiet wakefulness (Fig. 1B), while there was no significant difference in slow-wave sleep. Note that the cortical activity was often displaying Up/Down state dynamics in slow-wave sleep, and we performed an analysis (not shown), discarding the Down states prior to the response, and there was no significant change.

**Figure 1:**
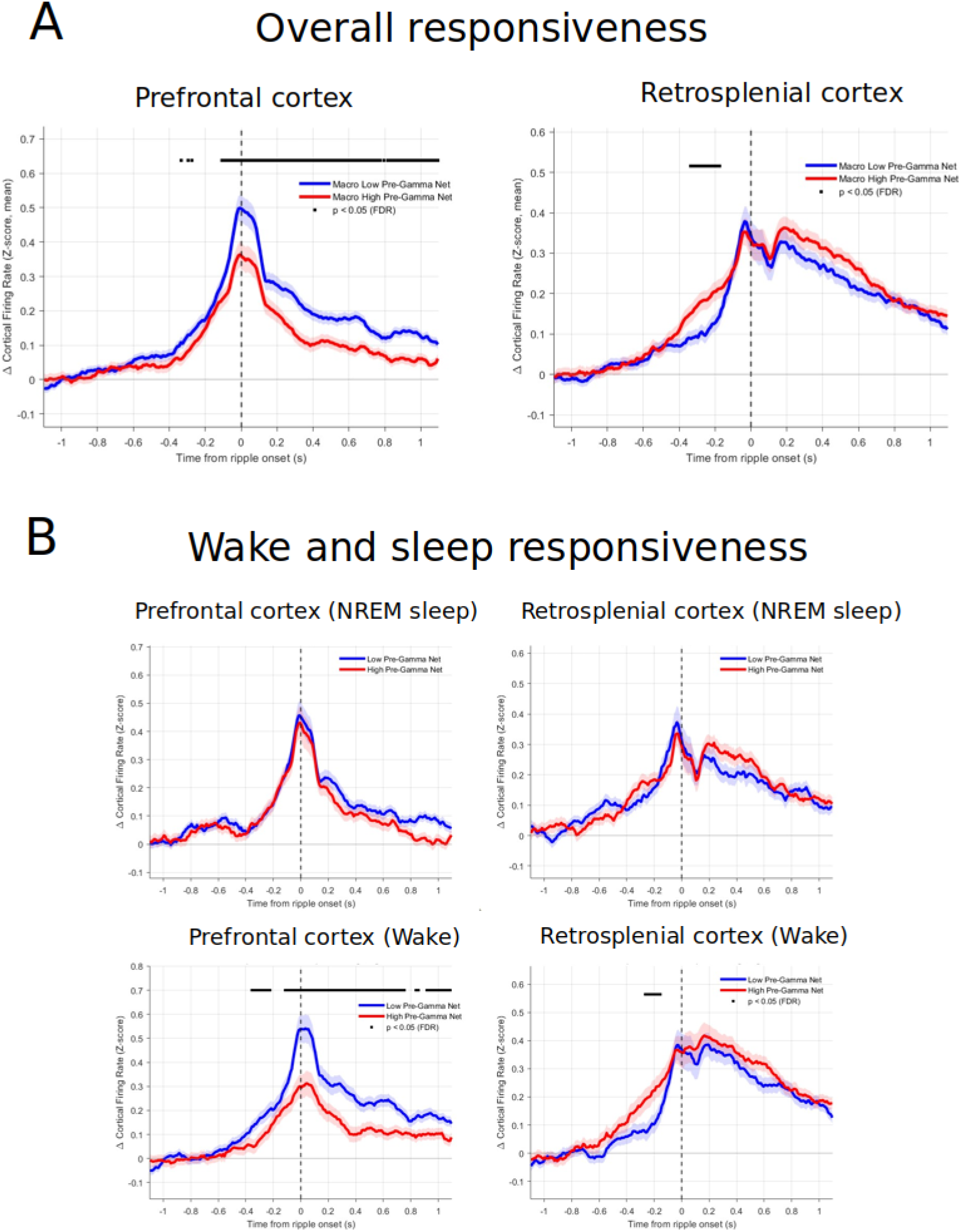
response in prefrontal and retrosplenial cortex to hippocampal sharp-wave ripple complexes. A. Overall spiking response in low-amplitude gamma oscillation states (blue) compared to high-amplitude gamma oscillations (red). B. Responses in the two brain areas were separately analyzed for wakefulness and slow-wave (NREM) sleep.

### 3.2 Computational models

To model this phenomenon, we used a previously introduced model of gamma oscillations (Tahvili & Destexhe, 2024), which is displayed in Fig. 2. A mean-field model was constructed for gamma oscillations, based on a spiking network model (Fig. 2A). The mean-field produced gamma oscillations for the same parameter range as the spiking model, and the frequency of the gamma oscillation was correctly captured by the mean-field (Tahvili & Destexhe, 2024).

**Figure 2:**
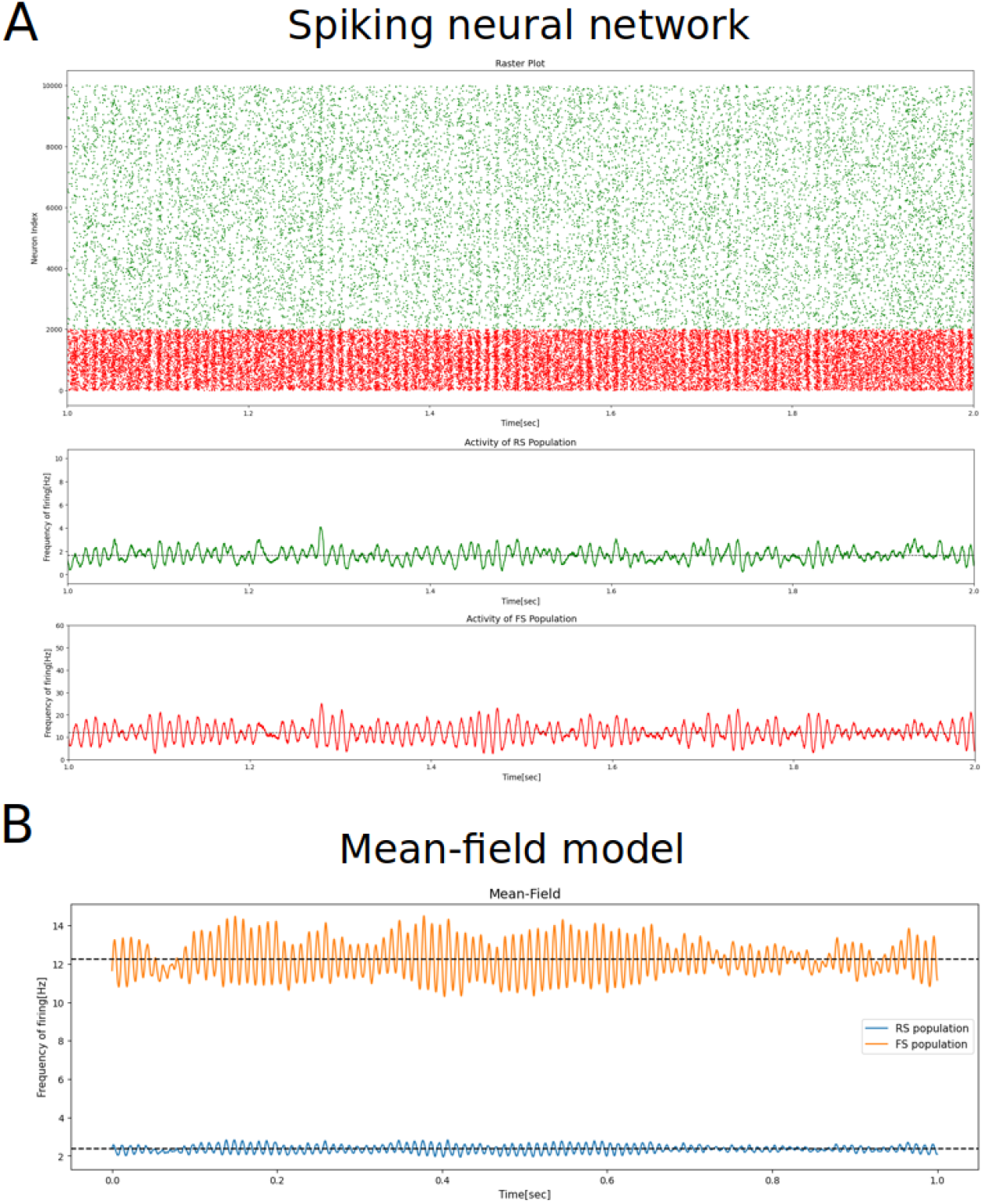
gamma oscillations in networks of excitatory (RS) and inhibitory (FS) neurons, comparing spiking networks (A) with the mean-field (B). The RS and FS neurons were displated in green and red, respectively, in A, while in B, the RS and FS populations were in blue and orange, respectively. Modified from Tahvili & Destexhe, 2024.

Using this model, we generated low- and high-amplitude gamma states by changing the resting level of RS cells (see Methods), with high-amplitude gamma oscillations were produced when RS cells are more depolarized. These two states are depicted in Fig. 3.

**Figure 3:**
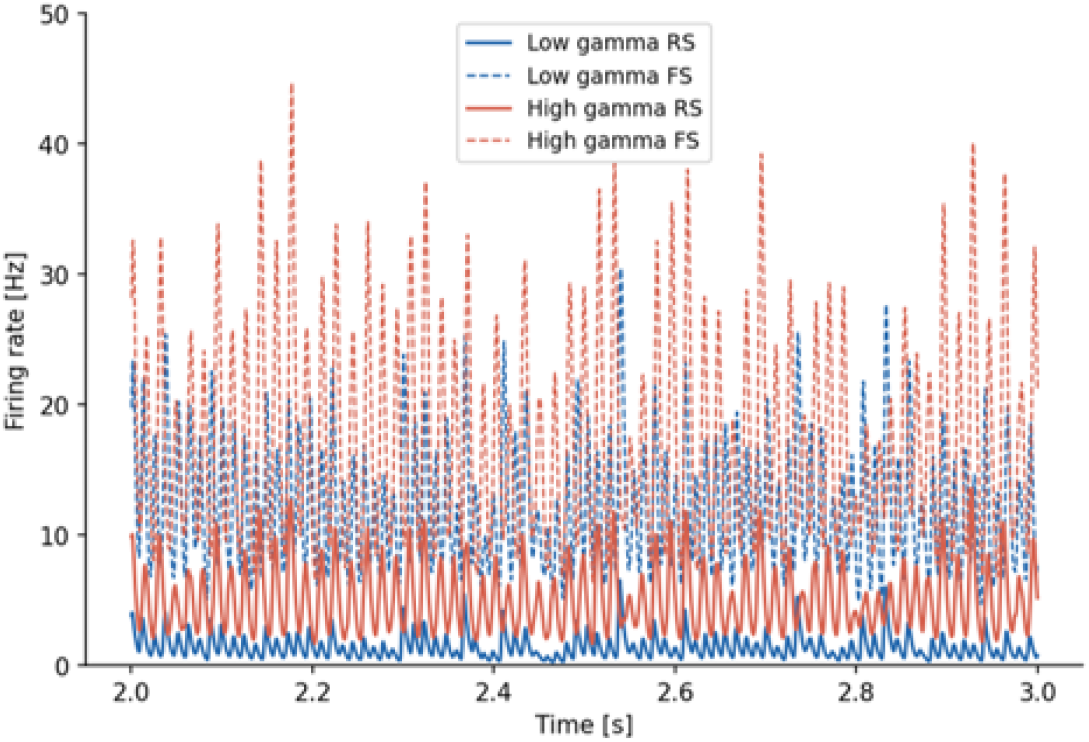
low-amplitude and high-amplitude gamma oscillations due to different resting membrane levels in RS cells (see Methods for parameter values).

Using these settings, we simulated an excitatory input equally distributed in the RS and FS cell populations. Figure 4 shows the response in the two gamma states, using a Gaussian-shaped input, with different temporal widths. We observe that in general the low-amplitude gamma state was more responsive compared to when the system exhibits high-amplitude gamma oscillations. Interestingly, this difference was essentially visible for RS cells, while FS cells showed no significant difference.

**Figure 4:**
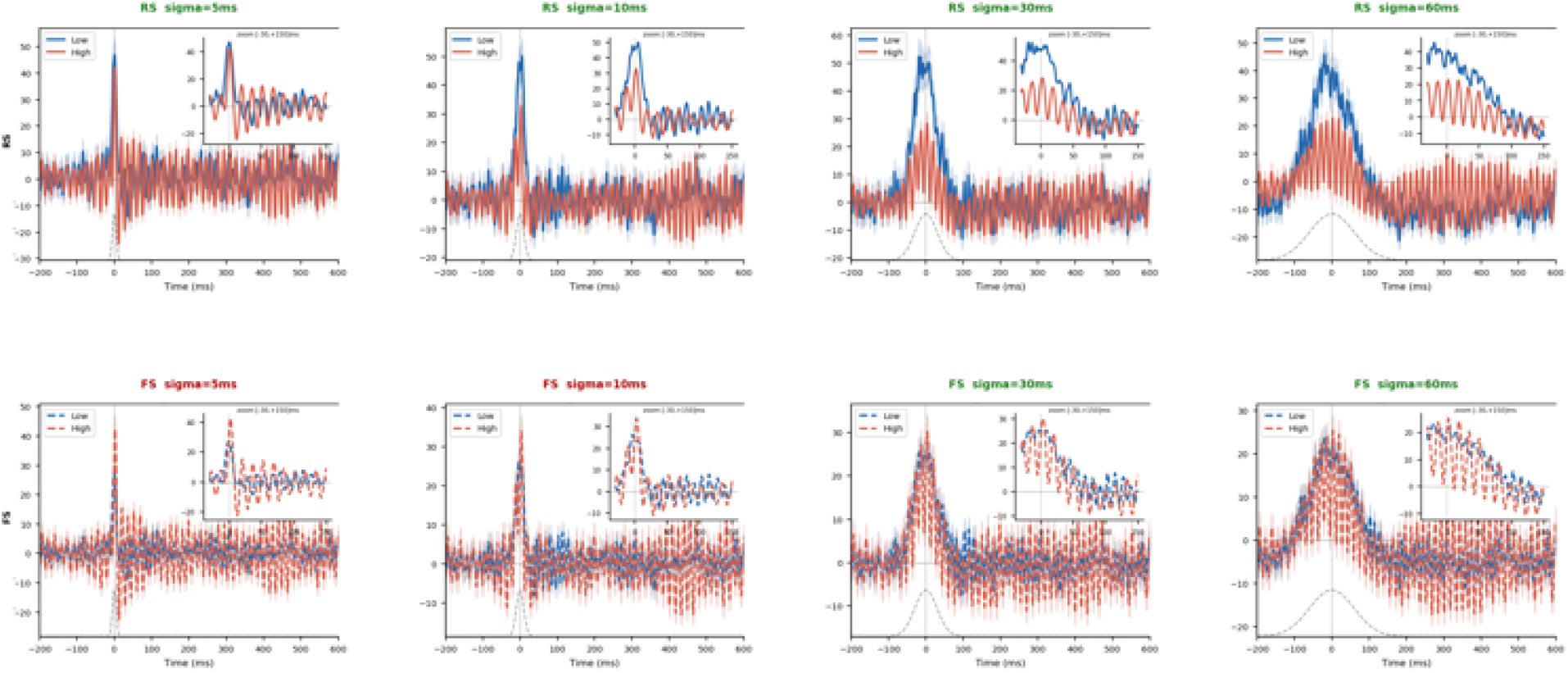
response of the mean-field model for low-amplitude (blue) and high-amplitude gamma oscillations. The response of RS cells are shown in the top panels, and FS cells in bottom panels. From left to right : different temporal width of the Gaussian shaped input.

We next checked the robustness of these findings with regard to recurrent excitatory connections. Figure 5 shows the response obtained for different levels of excitatory recurrence (excitatory-to-excitatory connections). We observe that the difference of responsiveness between low and high gamma states is robust against changes of excitatory connectivity. We also changed inhibitory connectivity and similarly, there was no significant change except for the absolute magnitude of the evoked response (not shown).

**Figure 5:**
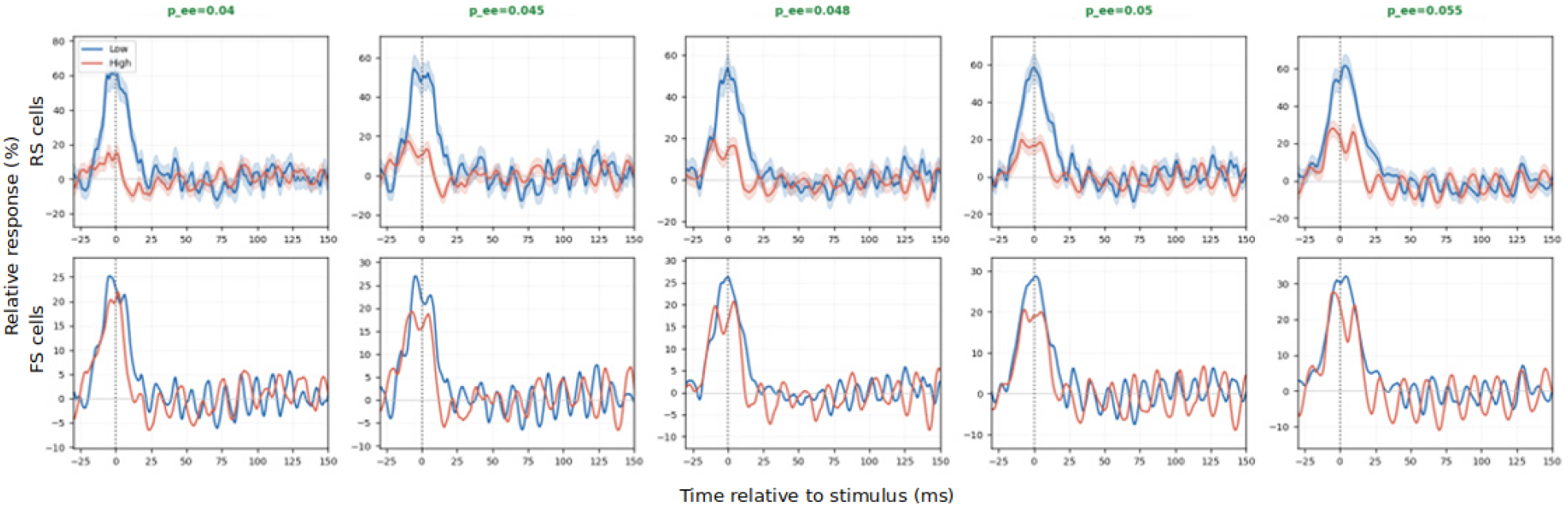
Robustness of the increased responsiveness in low-amplitude gamma states, for different levels of recurrent excitatory connectivity. The connection probability between RS cells is indicated on top.

We also evaluated the effect of the excitatory-inhibitory bias of the input. Figure 6 shows the response obtained for different inhibitory bias. Interestingly, one can see that when the input ratio was weaker on excitatory cells compared to inhibitory cells there was no difference between the gamma states (Fig. 6, left panels). Increasing the ratio in favor of excitatory cells resumed the difference of responsiveness (Fig. 6, right panels).

**Figure 6:**
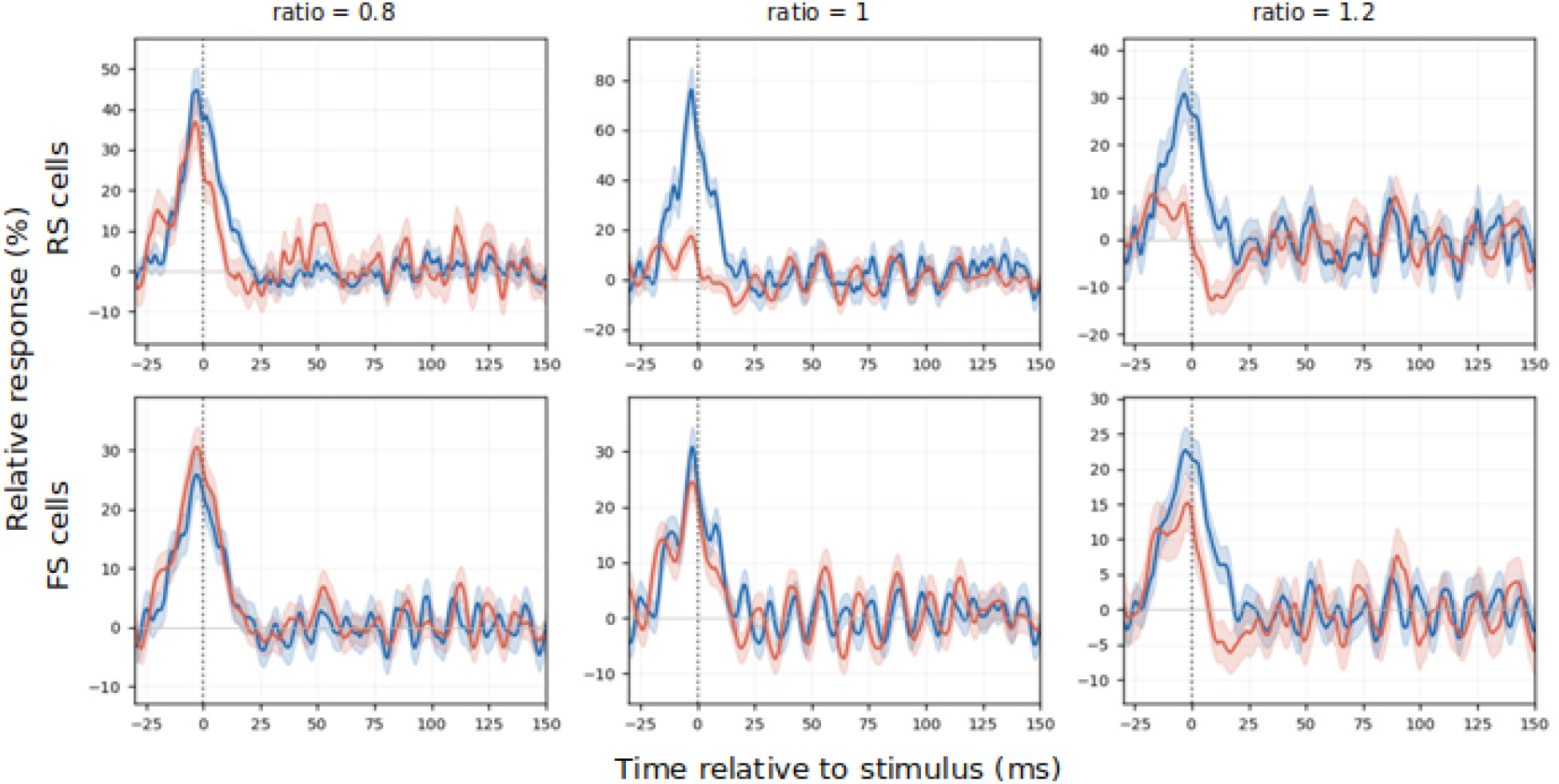
Responsiveness for different bias of excitatory and inhibitory input connections. The level of recurrent connectivity was lower (p=0.04), and the ratio indicates the proportion of connections to excitatory neurons. For a lower ratio (0.8), the evoked response was not significantly different between low-amplitude and high-amplitude gamma states.

We can see that for the vast majority of connectivity settings, the model reproduces the experimentally-observed difference of responsiveness in PFC. However, we also found a connectivity setting replicating the RSC observation of no significant difference (Fig. 6, left panels), using a reduced excitatory connectivity and a more important bias of the input connections towards inhibitory neurons.

Finally, we evaluated the consequences of such differences by simulating a chain of brain regions, each represented by one mean-field model. Figure 7 shows the input propagation resulting from this arrangement, as a function of the amplitude of gamma oscillations. As predicted by the differential response in Fig. 4, the inter-areal propagation was also dependent on the amplitude of gamma oscillations : for low-amplitude gamma states, the propagation of information was more reliable compared to high-amplitude gamma states.

**Figure 7:**
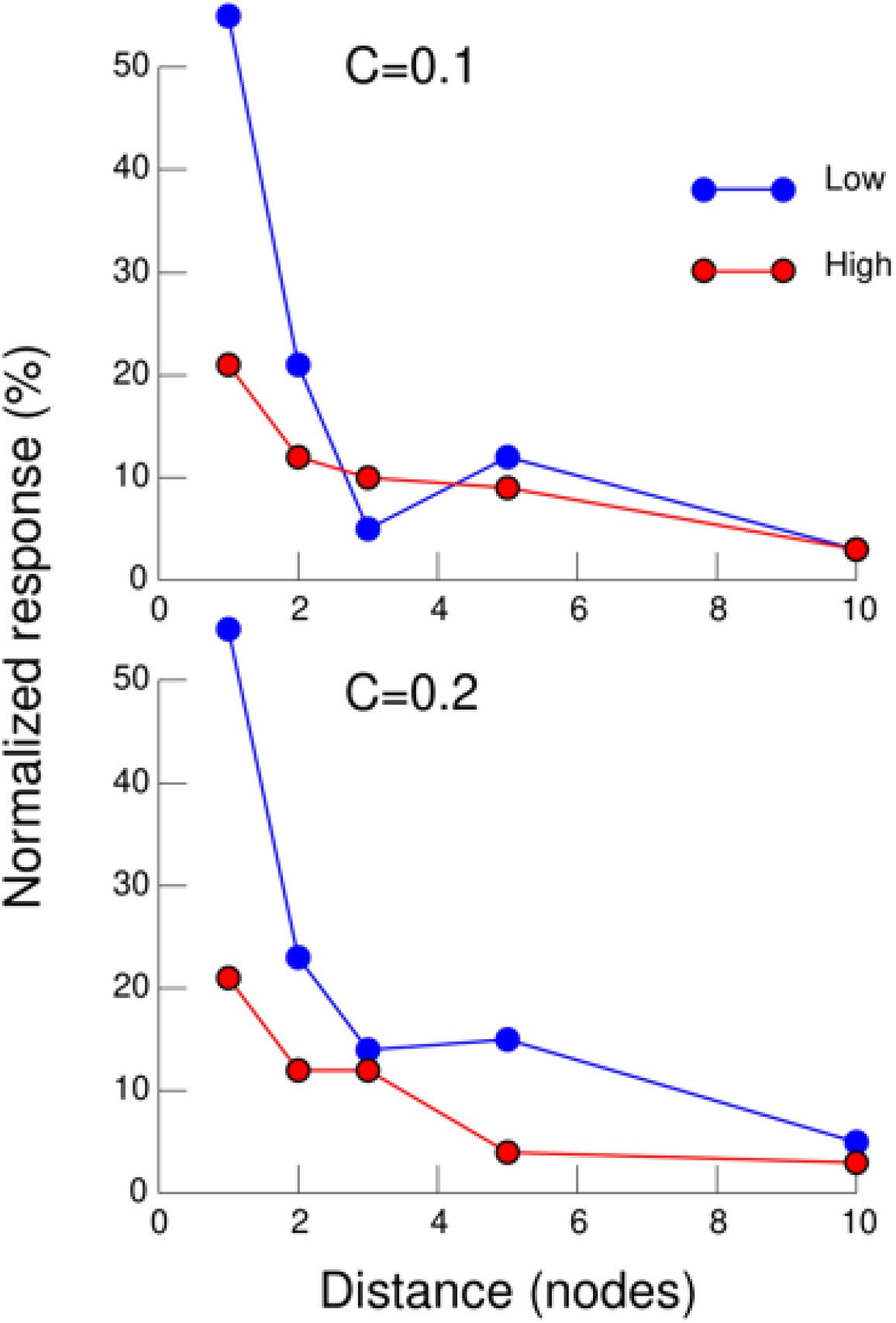
Prediction of the input propagation across different mean-field nodes, connected along a chain. The amplitude of the evoked response (normalized evoked firing rate) is shown here for two different coupling strengths. The input was given to node 1, and the response was measured in different nodes. The connectivity between nodes was first-neighbor and excitatory.

## Discussion

In this paper we have shown electrophysiological experiments on rats with simultaneous hippocampal and cortical recordings in two different anatomically-related regions, the CA1 region of hippocampus (dorsal and ventral) and two cortical regions, PFC and RSC. The main finding is that the cortical response depends on brain area and on brain state : in PFC, there is a significant difference between low-amplitude and high-amplitude gamma states, while no significant difference is seen in RSC. Using computational models, and in particular a mean-field model of cortical gamma oscillations (Tahvili & Destexhe, 2024), we could replicate the augmented responsiveness in low-amplitude gamma states. For particular connectivity settings, this difference vanished, which replicates the findings in RSC.

A first interesting observation is that the augmented responsiveness was found very robustly for variations of excitatory and inhibitory connectivity, as well as excitatory-inhibitory input bias. This suggests that there is indeed a better response, and better propagating activity over different connected regions (Fig. 5), when the cortical network exhibits none, or little gamma oscillations. This corroborates the previous finding of a diminished responsiveness of networks exhibiting gamma oscillations (Susin & Destexhe, 2021). In this regard, the experiments shown here constitute the first experimental evidence for such a diminished responsiveness in PFC.

A second main finding is that the RSC exhibits no difference according to the gamma state, which emphasizes that different brain areas can display different collective properties such as responsiveness. Here, the model could find such a non-significant difference, but only for reduced recurrent excitatory connectivity, and only if the input bias was favoring inhibitory cells (Fig. 6, left panels). There is indeed anatomical evidence that there is a reduced recurrent connectivity in RSC compared to PFC (Brennan et al., 2020) There is also some evidence that the input is biased towards inhibitory cells in RSC (Opalka et al., 2020) So this prediction from the model seems to be consistent with available anatomical data.

On a functional point of view, the prediction that there is an augmented inter-areal propagation in low-gamma states compared to high-amplitude gamma (Fig. 7) provides a functional interpretation about the role of gamma oscillations in cortex. In contrast to the communication through coherence hypothesis (Fries, 2005, 2015), our findings suggest that gamma oscillations would rather tend to isolate cortical regions from one another, favoring local computations. The absence of gamma would then be more favorable to large-scale communications between brain areas. This view offers the advantage of relieving the difficult constraint that gamma needs to be in phase between distant brain regions, in favor of a more flexible situation where information can be transmitted across large distances, regardless of the phase of the local network dynamics.

## Acknowledgments

Research supported by CNRS, the ANR (CR-CNS grant ImpactCom with NSF, FLAG-ERA grant BrainAct), the European Community (Virtual Brain Twin project, Horizon Health 101137289) and the ECOS-SUD France-Chile exchange program.

## Notes

### Competing Interest Statement

The authors have declared no competing interest.

